# Wall-following – phylogenetic context of an enhanced behaviour in stygomorphic *Sinocyclocheilus* (Cypriniformes: Cyprinidae) cavefishes

**DOI:** 10.1101/2023.08.31.555641

**Authors:** Bing Chen, Wen-Zhang Dai, Xiang-Lin Li, Ting-Ru Mao, Ye-Wei Liu, Marcio R. Pie, Jian Yang, Madhava Meegaskumbura

## Abstract

With 75 known species, the freshwater-fish genus *Sinocyclocheilus* is the largest cavefish radiation in the world, emerging as a model system for evolutionary studies. They show multiple adaptations for cave dwelling (stygomorphic adaptations), which include a range of traits such as eye degeneration (Normal-eyed, Micro-eyed and Eyeless), depigmentation of skin, and in some species, the presence of “horns”. Their behavioural adaptations to subterranean environments, however, are poorly understood. Wall-following (WF) behaviour, where an organism remains in close contact with the boundary demarcating its habitat when in the dark, is a peculiar behaviour observed in a wide range of animals and is enhanced in some cave dwellers. Hence, we hypothesize wall-following to be present also in *Sinocyclocheilus*, possibly enhanced in Eyeless species compared to species with visual cues (Normal / Micro-eyed species). Using 13 species representative of *Sinocyclocheilus* radiation and eye-morphs, we designed a series of assays, based on pre-existing methods for *Astyanax mexicanus* behavioural experiments, to examine wall-following behaviour under three stimulation conditions. Our results indicate that eyeless species exhibit significantly enhanced levels of WF compared to Normal-eyed species, with Micro-eyed forms demonstrating intermediate levels. Using a mtDNA based dated phylogeny (chronogram with four clades A – D), we traced the degree of WF of these forms to outline common patterns. We show that intensity of WF behaviour is high in the subterranean clades (B & C) compared to clades with free-living species (A & D). Experiments on WF behaviour revealed that eyeless species are highly sensitive to vibrations, whereas normal-eyed species are the least sensitive. Since WF behaviour is present to some degree in all *Sinocyclocheilus* species, and given that these fishes evolved in the late Miocene, we identify this behaviour as being ancestral with WF enhancement related to cave occupation. Our results from this diversification-scale study of cavefish behaviour suggest that enhanced wall-following behaviour may be a convergent trait across all stygomorphic cavefish lineages.

**Significance statement:** *Sinocyclocheilus*, a genus of 75 species of freshwater cavefish, is an emerging model system in evolutionary studies. Their adaptations for subterranean life, including eye degeneration, skin depigmentation, and horn-like structures, are well-known, but their behavioural adaptations remain understudied. Here we focus on a phenomenon, called “wall-following,” where fish stay close to the cave walls in absence of light. We hypothesized that this behaviour would be more pronounced in eyeless species. We selected 13 species, representative of the diversity of the genus and eye types, and observed their wall-following behaviour under different conditions. Results were intriguing; eyeless species exhibited heightened wall-following behaviour compared to their sighted counterparts, with small-eyed species falling in between. Researchers also mapped this behaviour on a phylogenetic tree, discovering a pattern: cave-dwelling clades showed stronger wall-following than free-living ones. Wall-following is prevalent in all *Sinocyclocheilus* species and, given the evolutionary history of the genus, is considered an ancestral behaviour that intensified with cave adaptation. These findings contribute to our understanding of convergent evolution, suggesting that enhanced wall-following may be a shared trait among diverse cavefish lineages.

## Introduction

Vertebrate lineages have evolved sensory systems and associated behaviours in order to adapt to new environments such as subterranean habitats. To occupy caves, species became adapted to the low availability of resources such as light, oxygen concentration and nutrients, leading to stygomorphic adaptations, including elongated appendages, lowered metabolism, specialized sensory systems, loss of eyes and pigmentation (Yoshizawa et al., 2012; Jeffery, 2019; Chen et al., 2020; Li et al., 2020; Ma et al., 2020a). A prominent stygomorphic convergent feature of cavefishes is the degeneration of eyes, compensated for by enhancements to the mechanosensory organs such as the neuromast lateral line system (Borowsky, 2013; Ma et al., 2020b; Chen et al., 2022b). A prominent swimming behaviour of cavefish is wall-following (WF, a form of thigmotaxis), where the fish senses the walls or boundaries of its cave environment in the absence of visual cues (Sharma et al., 2009; Patton et al., 2010). Although thigmotaxis has been reported in non-cavernicolous organisms introduced into a dark environment, this behaviour is putatively enhanced in cave-dwellers (Sharma et al., 2009; Norton, 2012; Niemiller and Soares, 2015).

Wall-following behaviour has previously been observed in freshwater-fish like *Astyanax mexicanus, Gasterosteus aculeatus* and *Danio rerio* (Patton et al., 2010; Johnson and Hamilton, 2017; Ginnaw et al., 2020). A lion’s share of this work has been on *A. mexicanus*, where some populations are cave-dwelling and exhibit distinct adaptations for cave life. These eyeless populations are capable of moving through complex environments without colliding with objects and their larvae prefer using a frontal approach using their head (Lloyd et al., 2018). Hence, cavefish resort to continuous swimming in order to constantly receive information from the environment (Windsor et al., 2008; Holbrook and Burt de Perera, 2013). Once cavefish detect a cave wall, they continue following the wall (Patton et al., 2010), which suggests that wall-following in cavefish is both spontaneous and continuous. This behaviour is enhanced in cavefish compared to other animals. Past studies have proposed wall-following behaviour as a strategy for foraging and spatial exploration (Sharma et al., 2009). In their perpetually dark environments, cavefish have evolved better short-range senses (hydrodynamic imaging ability) (Hassan, 1985; Windsor, 2014), such as tactile sensing using the anterior part of body (Sharma et al., 2009) and using mouth suction frequently to generate a suction flow to navigate non-visually (Holzman et al., 2014). This implies that wall-following might entails complex functions such as spatial orientation, seeking protection or refuge and obstacle avoidance.

Distinguishing between stationary and moving objects is of vital importance for cavefish due to the limited sensory perception in short distances within restricted cave environments. A study in *A. mexicanus* showed differences in WF behaviours under different boundary stimulations, observing that eyeless morphs swim nearly parallel to the wall, compared to sighted morphs. Under varying light conditions, eyeless morphs expressed closer swimming distance to the wall while reaching higher swimming speed compared to sighted morphs (Sharma et al., 2009). However, the evolution of wall-following behaviour (WF) in response to vision loss (morphological change) remains poorly understood.

With 75 species, *Sinocyclocheilus* genus (Cyprinidae, Barbinae) represents the largest cavefish radiation in the world (Jiang et al., 2019; Mao et al., 2021). These species show substantial morphological variability and inhabit suitable habitats of the massive 62,000 km^2^ south-western karstic landscape of China (Xiao et al., 2005; Romero et al., 2009; Zhao and Zhang, 2009; Jiang et al., 2019). They are phylogenetically well known, with four major clades (A – D), with clade B & C harbouring mostly the stygomorphic forms and clade A & D containing predominantly the surface dwelling forms (Mao et al., 2021). They represent an emerging model system for evolutionary novelty and show multiple adaptations for subterranean life.

For instance, they demonstrate varying degrees of eye degeneration, from Normal-eyed, to Micro-eyed and Eyeless (Meng et al., 2013; Zhao et al., 2021), loss of pigmentation (Li et al., 2020), absence of circadian rhythms, slow metabolism (Yang et al., 2016; Zheng, 2017), and a horn-like structure on the head on some species, of which the function is not clear (He et al., 2013). Studies on their natural history suggest that Eyeless cavefish are less active than Normal-eyed species. For instance, *S. grahami* (Normal-eyed, Surface-dwelling) swim faster and farther, swimming at a speed 2-3 times greater than *S. anshuiensis* (Zheng, 2017). The neuromast system in *Sinocyclocheilus* has been shown to be asymmetric and is correlated with the degree of eye-degeneration and is pronounced in the eyeless forms (Chen et al. 2022b). Furthermore, for several species, the wall-following behaviour is also thought to be associated with neuromast asymmetry, with eyeless forms having the strongest WF behaviours (Chen at al. 2022a). Deeper exploration is needed to understand the phylogenetic context and elucidate the range of stimuli influencing WF behaviour. Furthermore, a study of *Sinocyclocheilus* showed eyeless morphs showing being attracted in response to different stimuli (Chen et al., 2022a). Given the large number of species, the *Sinocyclocheilus* genus offers an ideal system for a deeper analysis of behaviour across a cavefish diversification.

Despite being an emerging multi-species model system, a radiation-scale understanding of *Sinocyclocheilus* cavefishes swimming behaviour is still lacking. Here, we investigate the swimming behaviour of *Sinocyclocheilus* species in a phylogenetic context. The species considered represent the three main habitat types and the three main eye-related morphologies: Normal-eyed (surface water bodies, Surface-dwelling habit), Micro-eyed (cave-associated habitats, Stygophilic habit) and Eyeless (cave habitat, Stygobitic habit) in the context of Mao et al. (2021). Given that WF behaviour is arguably enhanced in *A. mexicanus* cavefish populations and it is potentially affected by the sensory organs (e.g. neuromast in the lateral line system), we hypothesize that in *Sinocyclocheilus* species, WF is a shared, derived trait correlated with visual acuity – a characteristic for which we use eye-morphs as a proxy. Hence, we predict that Eyeless species will show the greatest level of WF behaviour, followed by Micro-eyed and Normal-eyed species, respectively. Furthermore, we predict that various (stable/ vibrative) stimuli will elicit a distinct response correlated to the extent of eye-degeneration condition.

## Materials & Methods

### Fish collection and maintenance

All experimental fishes (13 *Sinocyclocheilus* species, with 3 individuals from each species, N = 39,) were adults (Standard Length, SL: mean ± SD = 8.40 ± 1.28 cm) and collected from Yunnan and Guizhou Province and Guangxi Zhuang Autonomous Region of China between December, 2017 and September, 2020 (Supplementary Figure S1). Fish smaller than 8 cm were maintained in a Centralized Zebrafish aquarium system and housed in 1L BPA-free plastic tanks with each tank receiving separate water delivery and drainage. Fish larger than 8 cm were maintained in groups in four large aquariums (90 × 50 × 50 cm, 300 L; 150 × 80 × 80 cm, 1000 L capacity), equipped with dedicated filtration and purification equipment. Fish were regularly maintained on Shenyangkangcai™ fish food every day, consisting of shrimp, squid, spring fish and seaweed. We classified the species according to gross eye morphology as follows: Normal-eyed - *S. guilinensis, S. zhenfengensis, S. longibarbatus, S. macrophthalmus, S. oxycephalus, S. purpureus, S. maitianheensis*; Micro-eyed - *S. mashanensis, S. microphthalmus, S. bicornutus, S. multipunctatus*; Eyeless - *S. tianlinensis, S. tianeensis*. We designated the body shape as: fusiform - *S. macrophthalmus, S. oxycephalus* (Li, 1985; Wang and Chen, 1989); others were all categorized as compressiform (Zhao and Zhang, 2009) (Supplementary Table S1).

### Experimental equipment and video recording

For all assays, an individual was tested in one 45 × 28 × 28 cm rectangular assay arena. We used the aquarium system water (pH: 7.0 - 8.0, conductivity 150 - 300 S/m, temperature 19 ± 1°C, dissolved oxygen 8.5 mg/L), with water quality simulating natural conditions as much as possible. The depth of water was shallow (10 - 15 cm) depending on the size of fish, aiming to reduce their vertical excursions and to minimize depth of field errors with the movement tracking-system (as explained below). To minimize stress factors associated with difference in water properties, we changed the system water in tank after each assay. The fish were allowed a minimum of 10 minutes to acclimate and recover from the transfer process. Subsequently, the infrared illumination and digital video camera were activated. An infrared camera (Cannon XF 405) was set up about 1 m above the tank. An auxiliary infrared light source (850 nm; HongGuang, HG-IR1206, GangDong) uniformly irradiates the assay arena. We used a 4 Mbps (YCC 4:2:0, 25p, 1280 × 720) system setting to capture video under the infrared light (Figure 1A). Experimental design also referred to Chen et. al., 2022b. Due to our inability to visually track the cavefish in complete darkness, and in order to enhance the precision of software tracking, we adjusted our experimental settings. Noting that species with eyes ceased movement in total 0 Lux environments, all assays were conducted in a quiet, dimly lit room, simulating their natural habitat with a light intensity ranging from 1.7 to 5 Lux. We repeated the experiment three times with each fish. Except for when gravid, *Sinocyclocheilus* fishes do not show sexual dimorphism; together with given their rarity, we did not consider gender related behavioural differences. All individuals survived the experimentation.

**FIGURE 1.**
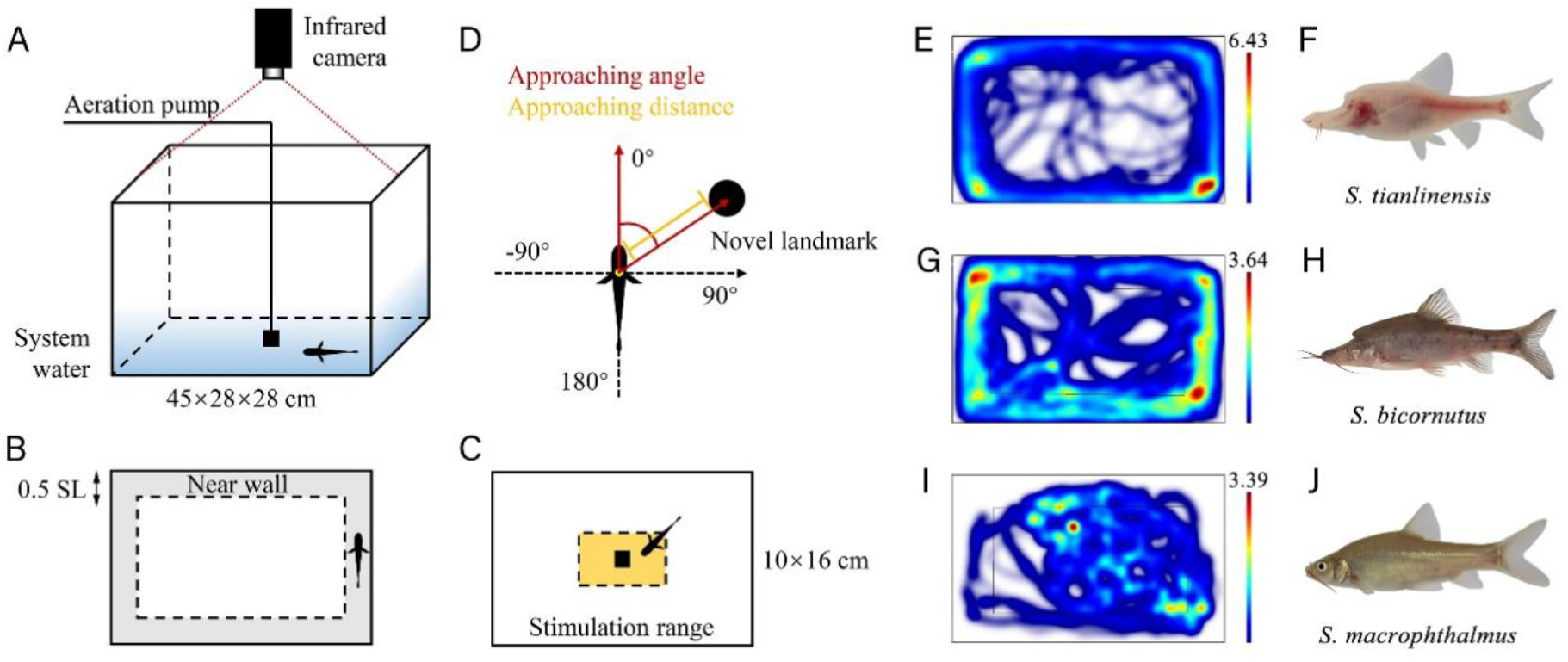
Diagram of the experimental apparatus and schematic diagram of measurements the representative trajectories of three species in the 10 minutes’ assay. Diagram of the experimental tank and equipment. (A) Vertical view of wall-following assay. The wall-following range is shown as the grey area, while its width is 0.5 SL. (B) Novel landmark and VAB assay’s stimulation range are shown in yellow. (C) Approaching angle and distance used for stimulation approaching analysis. (E,G,I) Representative swimming trajectories for the three forms: *S. tianlinensis* (Eyeless, Stygobitic, fusiform), *S. bicornutus* (Micro-eyed, Stygophilic, fusiform) and *S. macrophthalmus* (Normal-eyed, Surface, compressiform) in 10 minutes with no stimulation assay’s distance to the “near wall”. The colours represent the duration the fish spent in each pixel with low wavelengths (red) indicating greater times spent and high wavelengths (blue) indicating lower times spent at each pixel. Eyeless and Micro-eyed species maintain wall-following for a longer duration than the Normal-eyed species. Trajectory chart were created using the EthoVision XT software. (F,H,J) The three species for which the behaviour is depicted here.

### Wall-following assays

Since wall-following behavioural assays are established for *A. mexicanus* (Windsor et al., 2008; Sharma et al., 2009; Patton et al., 2010), we followed these in our study. We defined wall-following behaviour as swimming along the wall within a distance in 0.5SL and this area was called the near-wall belt (range of wall-following, Figure 1B). When fish maintained travelling a minimum distance for 2SLs within near-wall belt, we recorded it as wall-following behaviour. We used the EthoVision XT v.15 (Noldus IT, Wageningen, Netherlands) to track the swimming trajectories (Figure 2E, 2J and 2I). Standard length (SL) and pectoral fin length (PFL) were measured using FIJI (https://imagej.nih.gov) (Schneider et al., 2012). Swimming speeds less than 0.2 cm/s was set as immobility (resting) when analysing with EthoVision XT. We generated data for the following indicators in near-wall belt: WF-Distance (the distance fish swimming past within wall-following range), WF-Frequency (the frequency that fish swimming into wall-following range), WF-Time (the time that fish spend for wall-following), WF-Resting Time (the resting time during one assay), WF-Speed (the speed that fish swimming on average) and WF-Max Speed (the maximum speed fish reached during one assay; Table S8).

**FIGURE 2.**
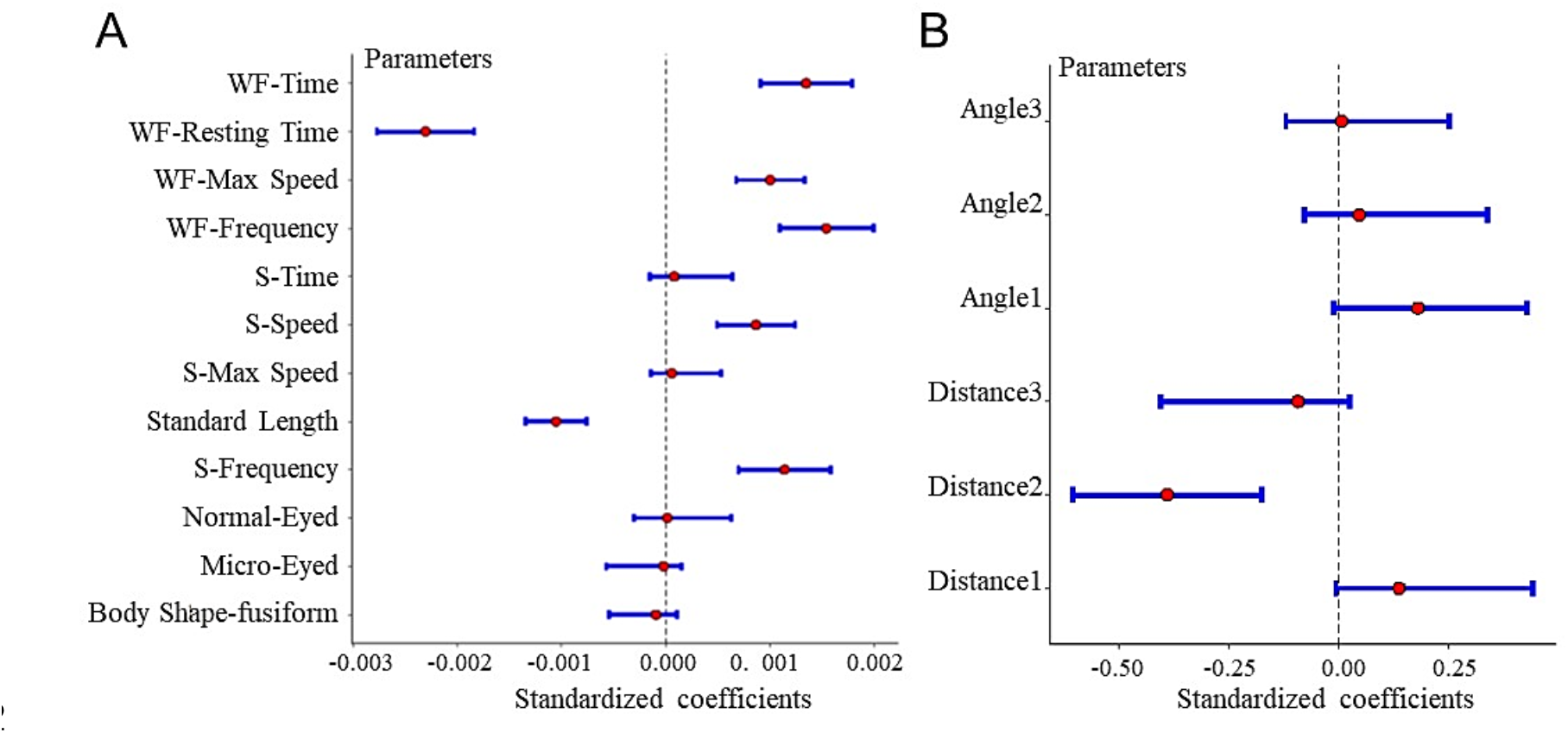
The effect of wall-following measurements parameters on the WF-Distance of assaying *Sinocyclocheilus* species (A) and the results of approaching angle/distance related with stimulation (B).

### Quantification of reaction to stimuli

To understand whether wall-following was a fixed behaviour or whether it could be affected by various stimuli, we performed three assays: one without stimulation, one with the novel-landmark and one with vibration attraction behaviour (VAB) setting. First, the unimpeded forward motion along the wall for 10 minutes was observed to figure out whether WF behaviour occurs (Video S1). Second, we tested for the novel-landmark for 5 minutes, by placing a dark opaque cylinder (diameter = 5cm) in the centre of rectangular arena, with the method referred to (Burt de Perera and Braithwaite, 2005; Lloyd et al., 2018) (Video S2). Third, we assayed vibration attraction stimulation for 3 minutes, with the method referred to (Jiang et al., 2016; Fernandes et al., 2018) (Video S3). Vibration was produced with an aeration pump (Jialu, LT-201S™, ZheJiang) working at 40 ∼ 50 Hz placed in the centre of the arena. The 10 × 16 cm rectangular area centred on stimulation was considered as the stimulation range (Figure 1C). We measured these indicators of fish swimming within this range: S-Frequency (the frequency fish swimming in), S-Time (the time fish spent in stimulation range), S-Speed (the speed fish swimming on average) and S-Max Speed (the maximum speed they reached during one assay).

We also measured the angle of approach (Approaching angle) and the distance of approach (Approaching distance) to stimulation by FIJI. Approaching distance was defined as the shortest distance between the edge of fish’s body and the stimulation (Figure 1D). Approaching angle was defined as the angle between a line extending down the midline of fish and a line extending from the centre of stimulation, both referred to Lloyd et al. (2018). In each assay, we recorded only the first three repeated approaches. Numbers from 1 to 3 represented the order of approaching within a given assay. For instance, ‘Angle 1’,’Distance 1’ indicated the first time the fish were attracted by the stimulation.

### Analysing the behaviour in an evolutionary context

We further analysed the evolution of wall-following behaviour from a phylogenetic perspective. We obtained two mtDNA fragments (Cytb and ND4) of 13 *Sinocyclocheilus* species and 5 outgroup species: *Cyprinus carpio, Puntius ticto, Labeo batesii, Gymnocypris przewalskii*, and *Gymnocypris eckloni* from Genbank (Supplementary Table S2). We edited and aligned sequences using Mega 6 (Tamura et al., 2013). BEAST v1.8.4 (Suchard et al., 2018) was implemented to build Bayesian time tree using two fossil calibration points (C1, C2) recommended by Li et al. (2008). Optimal substitution model and parameter settings were determined by jModeltest (Darriba et al., 2012). Previous studies have not yet provided definitive explanations regarding whether this behaviour is influenced by cave environments or if it is associated with ancestral cave preferences (Sharma et al., 2009; Patton et al., 2010). We then mapped behavioural patterns onto the inferred tree to assess their evolutionary trajectories.

### Statistical analysis

All the analyses were conducted in R v4.1.0 (R Foundation for Statistical Software, Vienna, Austria). Given that interspecific trait variation is often confounded by phylogenetic autocorrelation, traditional statistical methods might be subject to biases, such as elevated false-positive rates, thus requiring specific methods such as phylogenetic generalized least squares (PGLS) (Garamszegi, 2014) regression analysis. We tested for phylogenetic signal in different traits (WF-Distance, WF-Speed, SE-Diameter (eye trait, Table 1), WF-Frequency, WF-Time, WF-Max Speed), which was estimated using Pagel’s λ parameter in package “caper” (Freckleton et al., 2002; Orme et al., 2013). Result such as the value of λ nearing 1 denotes stronger signal (Pagel, 1999; Blomberg et al., 2003). Given that there was no evidence of phylogenetic signal in tested traits (λ = 0.000; Supplementary Table S4), we were justified in using non-phylogenetic statistical methods in all remaining analyses.

**TABLE 1.**
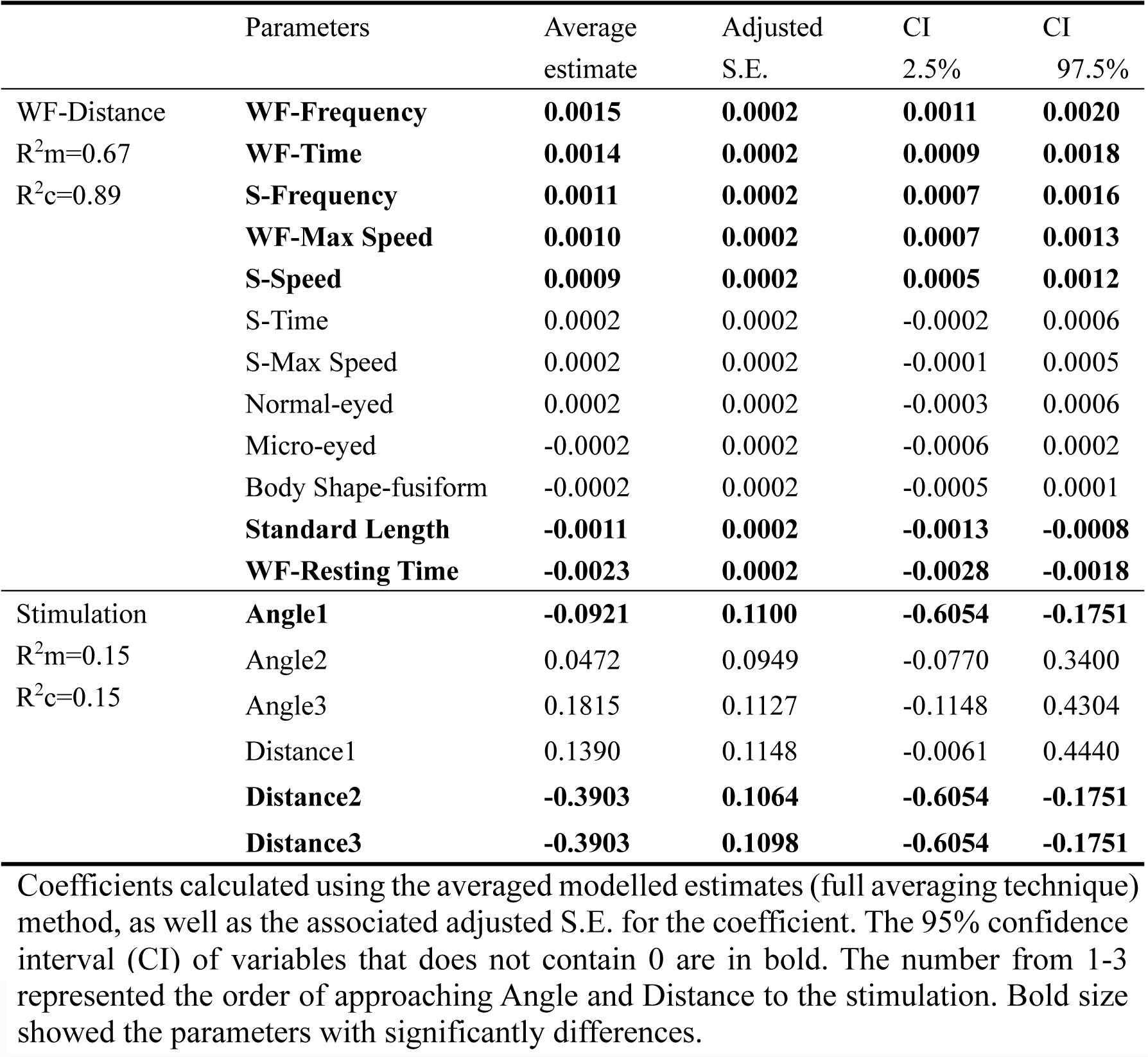
The results of model showing the variables that influenced WF-Distance and the approaching angle/distance related with stimulation.

|  | Parameters | Average estimate | Adjusted S.E. | CI 2.5% | CI 97.5% |
| --- | --- | --- | --- | --- | --- |
| WF-Distance<br>$R^2m=0.67$<br>$R^2c=0.89$ | **WF-Frequency** | **0.0015** | **0.0002** | **0.0011** | **0.0020** |
|  | **WF-Time** | **0.0014** | **0.0002** | **0.0009** | **0.0018** |
|  | **S-Frequency** | **0.0011** | **0.0002** | **0.0007** | **0.0016** |
|  | **WF-Max Speed** | **0.0010** | **0.0002** | **0.0007** | **0.0013** |
|  | **S-Speed** | **0.0009** | **0.0002** | **0.0005** | **0.0012** |
|  | S-Time | 0.0002 | 0.0002 | -0.0002 | 0.0006 |
|  | S-Max Speed | 0.0002 | 0.0002 | -0.0001 | 0.0005 |
|  | Normal-eyed | 0.0002 | 0.0002 | -0.0003 | 0.0006 |
|  | Micro-eyed | -0.0002 | 0.0002 | -0.0006 | 0.0002 |
|  | Body Shape-fusiform | -0.0002 | 0.0002 | -0.0005 | 0.0001 |
|  | **Standard Length** | **-0.0011** | **0.0002** | **-0.0013** | **-0.0008** |
|  | **WF-Resting Time** | **-0.0023** | **0.0002** | **-0.0028** | **-0.0018** |
| Stimulation<br>$R^2m=0.15$<br>$R^2c=0.15$ | **Angle1** | **-0.0921** | **0.1100** | **-0.6054** | **-0.1751** |
|  | Angle2 | 0.0472 | 0.0949 | -0.0770 | 0.3400 |
|  | Angle3 | 0.1815 | 0.1127 | -0.1148 | 0.4304 |
|  | Distance1 | 0.1390 | 0.1148 | -0.0061 | 0.4440 |
|  | **Distance2** | **-0.3903** | **0.1064** | **-0.6054** | **-0.1751** |
|  | **Distance3** | **-0.3903** | **0.1098** | **-0.6054** | **-0.1751** |

Coefficients calculated using the averaged modelled estimates (full averaging technique) method, as well as the associated adjusted S.E. for the coefficient. The 95% confidence interval (CI) of variables that does not contain 0 are in bold. The number from 1-3
represented the order of approaching Angle and Distance to the stimulation. Bold size showed the parameters with significantly differences.

We included the individual fish as a random effect to account for the repeated measures of each fish. To account for individual differences, WF-Distance and WF-Speed were expressed in terms of SL and WF-Time was calculated as the percentage of testing time (%). We formulated generalized linear mixed models (GLMM) to understand the effects of correlations and covariates in wall-following behaviour and stimulation approaching behaviour (two separate models) in 13 *Sinocyclocheilus* species by using the R package “lme4” (Bates et al., 2014; Nakagawa et al., 2017). As fixed independent variables, we used fish morphology (Eye-morphs, Body-shape, SL and PFL), wall-following measurements (WF-Frequency, WF-Time, WF-Resting Time, WF-Speed, WF-Max Speed) under: 10 minutes, 5 minutes, 3 minutes’ assays; and the stimulation range behaviour measurements (S-Frequency, S-Time, S-Speed, S-Max Speed). WF-Distance was set as response variable. Testing time was used as random independent variable. For model analysing approaching behaviour, we used approaching angle/distance as independent variables and stimulation as response variable. Since the data were diagnosed as over-dispersion, we selected negative binomial distributions for final models (Boswell, 1979). We used the package “MuMIn” to assess the AIC (Burnham and Anderson, 2002; Bartoń, 2019). We reported the fully-averaged results of the models as a ΔAICc threshold of 2. If the 95% confidence interval of a parameter is higher than zero, we considered it as an important factor explaining this model (Di Stefano, 2004; Nakagawa and Cuthill, 2007).

## Results

All *Sinocyclocheilus* cavefishes displayed a stereotypic “thigmotaxis” response to the new environment, revealed by an initial preference for wall-following of tank. Individual species exhibited variation in their responses to unfamiliar and new environments, such as sudden changes in motionlessness or swimming at a significantly faster pace. These variations possibly highlight the species-specific adaptations and behavioural strategies that come into play when encountering novel surroundings. However, the nature of wall-following behaviour differed between species, depending on the eye-morphs.

### Variables correlated with wall-following behaviour

WF-Frequency, WF-Time, S-Frequency, WF-Max Speed and S-Speed were the important variables affecting WF-Distance in *Sinocyclocheilus* species (Figure 2A, Table 1). Our results suggested that fish’s SL and WF-Resting Time were negatively correlated with WF-Distance; while WF-Frequency, WF-Max Speed, WF-Time, S-Frequency and S-Speed, were positively correlated with WF-Distance (Figure 2A). Our WF-distance and Stimulation model has R^2^_marginal_ value as 0.67 and 0.15, respectively. We checked the spatial autocorrelation and found none. Only the second approaching distance (Distance2) influenced stimulation negatively (Figure 2B, Table 1). We found that that all *Sinocyclocheilus* approached the stimulation at a narrow angle (< 90°; Supplementary Table S5). However, the approaching distance was significantly greater in surface fish than in cavefish (mean ± SD: Eyeless = 0.72 ± 0.91; Micro-eyed = 1.26 ± 1.07; Normal-eyed = 1.35 ± 1.08 SL), which suggested an ability to detect unknown objects at longer distances in Normal-eyed species.

### Wall-following ability is related with eye-morphs in *Sinocyclocheilus*

We found wall-following behaviour is pervasive in *Sinocyclocheilus* cavefishes, but with clear patterns associated with the eye-morphs. Considering studied here, except *S. purpureus*, more than 50% of time was spent following the wall (Figure 4, Time). Since Normal-eyed group spent the shortest WF-time, we found that eye regressed groups spent a longer in WF-Time (mean ± SD: Eyeless = 69.92 ± 21.76, Micro-eyed = 72.49 ± 14.58, Normal-eyed = 54.42 ± 24.39%, Kruskal-Wallis test: H_2_ = 27.55, *P* < 0.001; Supplementary Table S6). The average results of WF-Distance in Eyeless group were the longest, the shortest in Normal-eyed group and the Micro-eyed group was in between (mean ± SD: Eyeless 190.18 ± 128.74, Micro-eyed 156.20 ± 98.79, Normal-eyed 134.35 ± 105.14SL, Kruskal-Wallis test: H_2_ = 9.21, *P* < 0.05; Figure 3). Regarding WF-Frequency, Eyeless group wall followed most frequently, Normal-eyed group followed the least, with Micro-eyed group in between (mean ± SD: Eyeless = 24 ± 16, Micro-eyed = 22 ± 13, Normal-eyed = 18 ± 13 times, Kruskal-Wallis test: H_2_ = 8.62, *P* < 0.01). We also found that Eyeless species showed the highest WF-Speed, while Normal-eyed group swam at the lowest speed (mean ± SD: Eyeless = 0.52 ± 0.20, Micro-eyed = 0.45 ± 0.21, Normal-eyed = 0.36 ± 0.20SL/s, Kruskal-Wallis test: H_2_ = 22.77, *P* < 0.001). Since the Eyeless species showed the highest speed, the longest distance and the fastest speed in WF behaviour, they were considered as having the highest level of wall-following. Hence, our results show that WF behaviour in *Sinocyclocheilus* is enhanced with eye degeneration, with the highest enhancement in the Eyeless forms.

**FIGURE 3.**
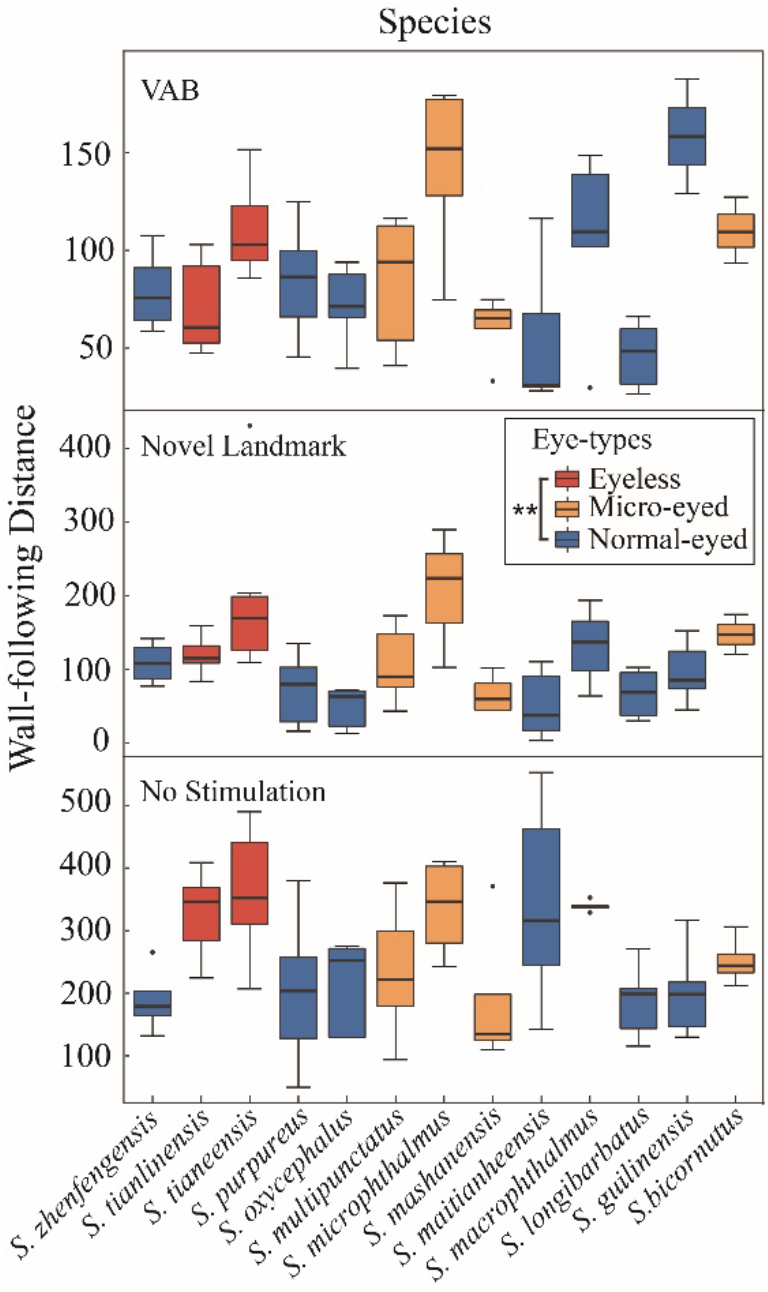
The results of wall-following distance of 13 *Sinocyclocheilus* species across three tests. 10 minutes with no stimulation, 5 minutes with Novel stimulation and 3 minutes with vibration attraction behaviour (VAB). Different colours on boxes indicate the Eyeless, Micro-eyed and Normal-eyed morphs. The wall-following distance in Eyeless groups were significantly higher than Normal-eyed groups (Kruskal-Wallis, Z=2.87, *P*=0.01). Wall-following distances are expressed in terms of standard length (SL). *: P < 0.05, **: P < 0.01, and ***: P < 0.001. Details of the statistical analyses are available in Supplementary Table S6.

**FIGURE 4.**
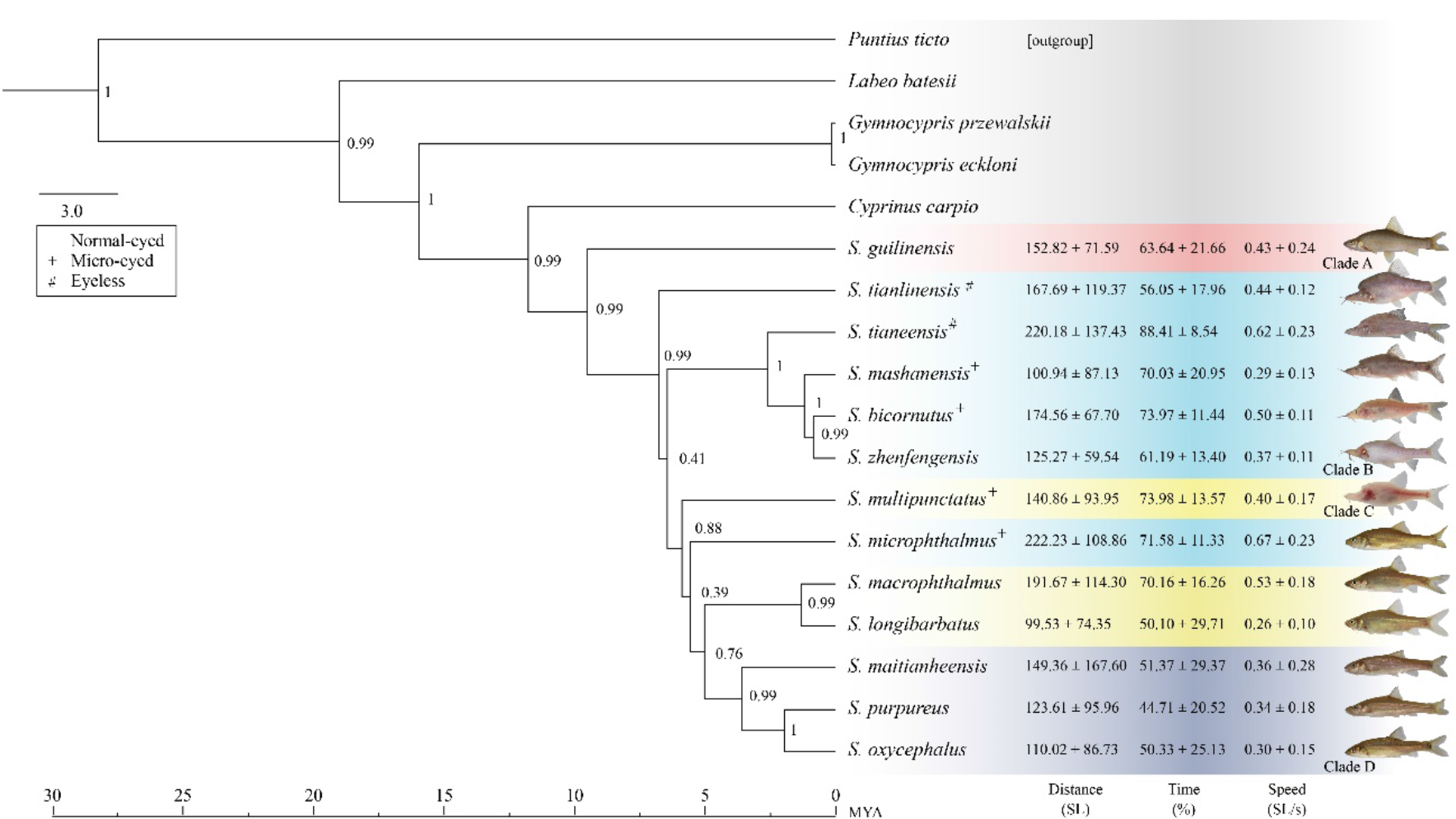
Bayesian inference tree derived from the concatenated data of mt-DNA and the mean results calculated from 10 minutes’ assay. Node values indicate clade posterior probability. Clade A-D are represented by four colours: Clade A – red, Clade B – blue, Clade C – yellow and Clade D – purple. The number represent the distance (SL), time (%) and speed of wall-following behaviour (SL/s), together with pictures of each species.

### Phylogenetic context of wall-following

The maximum credibility tree of *Sinocyclocheilus* shows four major clades (A – D), as previously reported by (Zhao and Zhang, 2009; Mao et al., 2021). The results suggest that wall-following behaviour is dominant in Clade B, where stygomorphism is greatest (the longest WF-Distance: *S. microphthalmus* = 222.23±108.86 SL, the longest WF-Time: *S. tianeensis* 88.41 ± 8.54 %, the highest WF-Speed *S. microphthalmus* 0.67±0.23 SL/s; showed in blue colour, Figure 4). We found that Clade B (mean ± S.D. = 172.37 ± 113.67 SL) showed significantly higher distance than Clade D (mean ± S.D. = 128.08 ± 119.75 SL; Dunnett’s T3 post-test: Z=3.50, *P*=0.05; Supplementary Table S7). Clade B also showed the fasted WF-Speed, the longest WF-Time and the most WF-Frequency compared with other clades (Supplementary Table S7).

However, from Clade C to D, with a predominance of normal sighted species, wall-following behaviours decreased (the shortest WF-Distance: *S. longibarbatus* 99.53±74.35 SL, the shortest WF-Time: *S. purpureus* 44.71±20.52 %, the lowest WF-Speed: *S. longibarbatus* 0.26±0.10 SL/s; Figure 4). But, we found one species to be an outlier to this pattern – *S. macrophthalmus* (Normal-eyed, Stygobitic, Clade C) has a great WF-Distance as long as the Eyeless cavefishes.

## Discussion

*Sinocyclocheilus* diversification began with the advent of the polar ice caps and the rain-shadow effect of the Himalayas, as a result of which the Guangxi, Guizhou and Yunnan region became drier in the late Miocene (Mao et al., 2021). This provided time for diversification across this vast karstic landscape, where species acquired, to different degrees, depending on the habitat, many stygomorphic traits. Our study contributes to this unfolding understanding by revealing wall-following behaviour as a key evolutionary adaptation within this genus. This behaviour, also appearing in unrelated lineages such as *A. mexicanus* cavefish, paves the way for a deeper exploration of the forces driving this convergent evolution. Therefore, it is crucial to consider the variability within and among clades, and the influence of ecological and evolutionary factors on wall-following behaviour.

Our analysis showed that wall-following level was the highest in Eyeless species and the lowest in Normal-eyed species. We mainly evaluated three aspects of wall-following level in *Sinocyclocheilus* genus: time, speed and distance with being the most important, as distance is a product of speed and time. However, we further considered speed and time also, for they can explain more subtle aspects of behaviour (Hoke et al., 2012). We noted the enhancement in wall-following ability going from Normal-eyed to Eyeless species. Eyeless species exhibited the most enhanced wall-following behaviour (highest speed, distance and a longer time in WF behaviour), however, it is not consistently supported across all measured variables (such as WF-Time and WF-Max Speed). Though our statistical analyses (Supplementary Table S6), confirmed a nonsignificant gradient from Eyeless to Normal-eyed species for these variables. In our analysis, we retained extreme results, not excluding outliers, as these data points could reveal interesting insights for future in-depth analyses involving more taxa. The variations we observed in the data can be attributed to individual differences among species. For instance, the Micro-eyed species (*S. microphthalmus*) exhibited a range comparable to some Eyeless species, further highlighting the complexity and diversity within the studied group. Moreover, the term “enhance” does not necessarily imply a linear relationship between the degree of eye development and the magnitude of wall-following behaviour. It reflects, instead, a behavioural divergence potentially influenced by various factors including sensory capabilities, ecological pressures, and phylogenetic history. As such, the proposition that Micro-eyed species represent an “in-between” state in terms of wall-following behaviour variables is not strongly supported by our data and requires further scrutiny.

The experiments were conducted under near-dark conditions, as observing and recording cavefish behaviour proved challenging in zero lux conditions. Additionally, it was noted while caring for the fish that in absolute darkness, the cavefish ceased movement. Interestingly, most of the cave entrances from which these cavefish were sampled were not in complete darkness, though some species were captured from completely dark environments well within caves. Therefore, the light values applied in our experiments corresponded with the natural low-light conditions observed in the field.

The cave habitats are low in terms of resources and predators (Romero et al., 2009; Ajemian et al., 2015; Niemiller and Soares, 2015). Past studies pointed out that swimming behaviour is suggested as being important in exploration of habitats (Teyke, 1985). In addition, wall-following is thought to be an extension of the swimming for exploration (Sharma et al., 2009). The faster swimming speed of cavefish enable fish to acquire more information via amplitude of water flow (Teyke, 1988) and to explore the cave environment continuously. Our findings on wall-following behaviour is also suggestive of different strategies for resource utilization and risk avoidance. Continuous swimming behaviour associated with wall-following, as seen in Eyeless species, may have evolved to optimize resource utilization in the resource-poor, predator-scarce cave environment. Continuous wall-following has already been shown in *A. mexicanus* (Patton et al., 2010; Yoshizawa, 2015) and has been explained in terms of exploratory spatial awareness. Hence, continuous swimming behaviour associated with wall-following may have evolved to enable fish to acquire more information via the amplitude of water flow, explore their surroundings continuously, so as to optimize resource utilization in the near absence of predation. On the contrary, the slower WF-speed in Normal-eyed species might reflect a defensive strategy to reduce the risk of injury. This dual role of wall-following behaviour in exploration and defence underscores its ecological and evolutionary significance. In other groups, it has been shown that the narrow and fixed wall-following route could be used as escape routes should threats arise (Sharma, 2008; Ajemian et al., 2015; Ginnaw et al., 2020), while preserving energy to accelerate rapidly when the need arises.

Eyeless species approached stimulation at a narrow angle, while surface-living species showing a greater approaching distance than cavefish. This could indicate an ability to detect unknown objects at longer distances in Normal-eyed species, reflecting differences in sensory capabilities and risk assessment strategies among the species. This spatial exploratory behaviour also associated with the enhanced olfactory system and lateral line systems in cavefishes (Fernandes et al., 2018; Kasumyan and Marusov, 2018; Lloyd et al., 2018; Chen et al., 2022b).

Some studies on other lineages about wall-following behaviour highlight different functional explanations, such as risk-avoidance in the three-spined sticklebacks (Ginnaw et al., 2020) and also mainly as an exploratory strategy on Somalian cavefish *Phreatichthys andruzzii* (Sovrano et al., 2018). The possible reason is that once the behaviour was changed as an adaptation to cave environments, it is more likely to persist under relaxed selection (Hoke et al., 2012).

Our results also offer some support for the enhancement of wall-following behaviour in stygomorphic species, particularly those in Clade B, the patterns are not uniformly clear-cut across all variables or clades. The intricate interplay of genetics, development, ecology, and environment shapes the evolution of behavioural traits in these species. The lack of strong phylogenetic correlation in the tested eye-morphs and behavioural traits in our study may be due to inadequate taxon sampling, the inclusion of species from only one of the clades that contain Eyeless species (Clade B). Due to rarity and sampling related problems, we could not analyse wall-following behaviour of stygomorphic species from Clade D, such as *S. anophthalmus*. However, the species from Clade B, *S. tianlinensis* and *S. tianeensis* share a common ancestor and thus supports the evolution of Eyeless species related intense wall-following behaviour due to phylogenetic inertia. Moreover, the evolutionary convergence of wall-following behaviour in unrelated lineages, as observed in our study, supports the idea that this behaviour has evolved in response to similar selective pressures. The prevalence of wall-following to various degrees across the phylogeny suggest that the trait is ancient.

To summarize, our study provides a complex, multi-faceted picture of wall-following behaviour in *Sinocyclocheilus* species and its relationship with eye morphology and phylogenetic clades. Low intraspecific variability of wall-following suggested that this behaviour was fixed for a species. Some comparable results from other cavefish lineage, such as in *A. mexicanus* cavefishes, show the similar wall-following swimming patterns (Patton et al., 2010). Evolutionary convergence of wall-following support the idea that this behaviour has evolved in response to similar selective pressures in evolutionarily unrelated lineages. The prevalence of wall-following, to various degrees across the phylogeny suggest that the trait is ancient and shared in cavefishes. The insights gained from such research can shed light on the broader patterns and processes of evolution in cave-dwelling organisms. Future research should continue to explore these relationships, taking into account the inherent variability and complexity of behavioural traits in these fascinating taxon.

## Conclusion

Our diversification scale behavioural assays show that *Sinocyclocheilus* have wall-following behaviours associated with cave-dwelling propensity. The Eyeless species showed the highest level of wall-following behaviour and Normal-eyed showed the least level, with Micro-eyed forms in-between. Our study confirmed that wall-following is correlated with multiple factors, especially with wall-following frequency, time and eye-morphs. Though the determination of the exact function of wall-following needs further experimentation, we suggest that wall-following facilitate protection and foraging behaviour in Eyeless forms, and for defence in eyed species. We found that wall-following is enhanced in Clade B and C (regressed-eyed species) but reduced in Clade A and D (Normal-eyed species). However, our results do not show phylogenetic correlation of wall-following behaviour, possibly due to inadequate taxon sampling. The convergence of wall-following with *A. mexicanus* cavefish suggest that this behaviour is an adaptation in response to selective regimes of subterranean environments. Our work will also form the foundation for further behavioural work on this emerging multi-species evolutionary model system.

## Abbreviations

WF: Wall-following
S: stimulation
VAB: vibration attraction behaviour
SL: standard length
PFL: pectoral fin length
GLMM: generalized linear mixed models
AIC: Akaike Information Criteria
PGLS: phylogenetic generalized least squares analysis
λ: lambda.

## Data availability statement

The raw data supporting the conclusions of this article will be made available by the authors, after the publication of the peer-reviewed paper.

## Ethics statement

Inspection of this study approved by the Guangxi University (GXU-2021-125; 2022-GXU-005). The study was approved by the National Natural Science Foundation of China and the Innovation Project of Guangxi Graduate Education. This study conforms with the regulation of the People’s Republic of China and the ASAB/ABS Guidelines for the Use of Animals. The fish were captured using umbrella nets, transported in oxygenated plastic bags with water from the cave habitat inside cooler box (18 - 20°C). In captivity, the cave habitat was mimicked in terms of water hardness, temperature and absence of light (for the cave species). We observed the cavefish without any harm to them or any other treatments with chemicals.

## Author contributions

Conceptualization: BC, MM, JY; Investigation: BC, MM, XL-L, TR-M, YW-L, MP, JY; Methodology: BC, MM, TR-M, MP, JY; Project administration: MM; Resources: MM, JY; Supervision: MM, MP, JY; Validation: BC, MM, WZ-D, TR-M, YW-L, JY; Visualization: BC, WZ-D, TR-M, YW-L, JY; Roles/Writing – original draft: BC, MM, WZ-D; Data curation: BC, XL-L, YW-L, JY; Formal analysis: BC, WZ-D, XL-L, TR-M, MP; Funding acquisition: MM, JY. All authors contributed to the article and approved the submitted version.

## Funding

This work was supported by the (1) Startup funding for MM though Guangxi University. (2) National Natural Science Foundation of China (#32260333) to MM. (3) National Natural Science Foundation of China (#31860600) to JY for fieldwork. (4) Innovation Project of Guangxi Graduate Education (#YCBZ2021008) to TRM and CB for research work. These funding bodies played no role in the design of the study and collection, analysis, and interpretation of data or in the writing of the manuscript.

## Conflict of Interest Statement

The authors declare that the research was conducted in the absence of any commercial or financial relationships that could be construed as a potential conflict of interest.

## Acknowledgments

We thank the EED lab members for fieldwork; Cheng-Hai Fu for the field working; Rohan Pethiyagoda (Ichthyology Section, Australian Museum) commenting on the manuscript.

## Notes

### Competing Interest Statement

The authors have declared no competing interest.

